# Base pairing interactions between substrate RNA and H/ACA guide RNA modulate the kinetics of pseudouridylation, but not the affinity of substrate binding by H/ACA small nucleolar Ribonucleoproteins

**DOI:** 10.1101/569061

**Authors:** Erin Katelyn Kelly, Dominic Philip Czekay, Ute Kothe

## Abstract

H/ACA small nucleolar ribonucleoproteins (snoRNPs) pseudouridylate RNA in eukaryotes and archaea. They target many RNAs site-specifically through base-pairing interactions between H/ACA guide and substrate RNA. Besides ribosomal RNA (rRNA) and small nuclear RNA (snRNA), H/ACA snoRNPs are thought to also modify messenger RNA (mRNA) with potential impacts on gene expression. However, the base-pairing between known target RNAs and H/ACA guide RNAs varies widely in nature, and therefore the rules governing substrate RNA selection are still not fully understood. To provide quantitative insight into substrate RNA recognition, we systematically altered the sequence of a substrate RNA target by the *Saccharomyces cerevisiae* H/ACA guide RNA snR34. Time courses measuring pseudouridine formation revealed a gradual decrease in the initial velocity of pseudouridylation upon reducing the number of base pairs between substrate and guide RNA. Changing or inserting nucleotides close to the target uridine severely impairs pseudouridine formation. Interestingly, filter binding experiments show that all substrate RNA variants bind to H/ACA snoRNPs with nanomolar affinity. Next, we showed that binding of inactive, near-cognate RNAs to H/ACA snoRNPs does not inhibit their activity for cognate RNAs, presumably because near-cognate RNAs dissociate rapidly. We discuss that the modulation of initial velocities by the base pairing strength might affect the order and efficiency of pseudouridylation in rRNA during ribosome biogenesis. Moreover, the binding of H/ACA snoRNPs to near-cognate RNAs may be a mechanism to search for cognate target sites. Together, our data provide critical information to aid in the prediction of productive H/ACA guide – substrate RNA pairs.

## Introduction

H/ACA snoRNAs constitute a highly versatile machinery that introduces pseudouridine modifications in rRNA and snRNA, but most likely also in mRNA and long non-coding RNAs (Carlile et al. 2014, Schwartz et al. 2014, Yu and Meier 2014). Pseudouridines are both the most abundant, yet also the subtlest modifications of cellular RNAs as they are C-C glycosidic isomers of uridine (Spenkuch et al. 2014). The entire functional impact of pseudouridines in cellular RNA is not yet fully understood, but pseudouridines have been shown to enhance RNA stability, ribosome function, as well as branch-site interactions in the spliceosome, and pseudouridines in mRNA have been speculated to regulate gene expression (Arnez and Steitz 1994, Yang et al. 2005, Liang et al. 2009, Carlile et al. 2014, Schwartz et al. 2014). The site-specific formation of pseudouridines is catalyzed either by H/ACA snoRNPs found in archaea and eukaryotes or by stand-alone pseudouridine synthases present in all domains of life (Rintala-Dempsey and Kothe 2017). H/ACA snoRNPs are particularly versatile as they are able to isomerize a large number of specific uridines to pseudouridines by utilizing a number of different H/ACA guide RNAs that recognize target RNA by base-pairing interactions (Ganot et al. 1997, Ni et al. 1997). H/ACA snoRNPs are composed of one H/ACA guide RNA bound to four conserved proteins (Cbf5/dyskerin, Nop10, Gar1, Nhp2 in yeast/humans) of which Cbf5/dyskerin is the catalytic entity. Notably, H/ACA guide RNAs share a conserved structure harboring typically two hairpins with an internal loop called the pseudouridylation pocket (Ganot et al. 1997). The target RNA base-pairs in a bipartite fashion to both sides of the pseudouridylation pocket leaving the target uridine and typically one additional nucleotide on the 3′ side of the target uridine unpaired at the base of the upper stem of H/ACA guide RNA. Besides the common structure, H/ACA guide RNAs are characterized by two short sequence motifs, the H box with the sequence ANANNA located in the hinge region between the two hairpins and the ACA box following the second hairpin at the 3′ end of the H/ACA guide RNA. Otherwise, H/ACA guide RNAs are highly diverse in their sequence and in their interactions with their respective substrate RNAs rendering it difficult to predict active pairs of H/ACA guide RNAs and target RNAs with pseudouridines.

The structure and stability of the H/ACA guide RNA is critical for efficient substrate RNA binding and pseudouridylation. Previously, it was determined that not all predicted guide RNA-target RNA pairs resulted in successful pseudouridylation. The modification ability is strongly dependent on three factors, namely the stability of the guide RNA hairpin, sufficient base pairing between the guide RNA and the target RNA in the pseudouridylation pocket, and a conserved distance between the target uridine and the H or ACA boxes (Xiao et al. 2009). The stability of the guide RNA hairpins contribute to substrate RNA binding through coaxial stacking interactions that form between both the upper and lower stems of the H/ACA guide RNA and the new helices formed by the binding of the substrate RNA to the single stranded pseudouridylation pocket (Liang et al. 2007, Duan et al. 2009). The conserved distance of 14-16 nucleotides between the target uridine and the H or ACA boxes is important for properly aligning the substrate to the catalytic domain of Cbf5, such that the target uridine can be appropriately docked into the active site (Wu and Feigon 2007, Duan et al. 2009, Caton et al. 2018). The interaction between the H/ACA guide RNA and the substrate RNA has been characterized as an omega structure, forming a three-way junction between the two guide-substrate RNA helices on either side of the pseudouridylation pocket and the upper stem of the guide RNA (Jin et al. 2007, Wu and Feigon 2007). The substrate RNA only interacts with the guide RNA on one side, rather than being threaded through the pseudouridylation pocket, and the H/ACA proteins assemble on the opposite side of the guide RNA.

The interactions between H/ACA guide RNAs and their natural substrate RNAs is best understood in *Saccharomyces cerevisiae* where all known H/ACA guide RNAs except one (snR30) modify known sites in rRNA or snRNA (Torchet et al. 2005, Piekna-Przybylska et al. 2008). In contrast, many human H/ACA guide RNAs are labelled as orphan since no corresponding target RNAs are known so far. In yeast, the nature of the base pairing between the guide RNA and the substrate RNA varies dramatically between H/ACA guide-target RNA pairs. There is no consistent length of base pairing region on either the 5□ or 3□ side of the pseudouridylation pocket, and an inconsistent number of non-canonical pairs and mismatches can be accommodated in the pseudouridylation pocket. For example, the snR191 3□ hairpin makes only 8 base pairs with its substrate in the 25S rRNA, with 4 base pairs formed on either side of the target uridine. The longest known interaction in nature is 17 base pairs, and it occurs between the 3□ hairpin of snR82 and the 25S rRNA. The fewest number of base pairs made on one side of the target uridine is 3 base pairs (e.g. in snR3 and snR81), and the maximum number of base pairs on one side is 10 base pairs in snR82. Typically, the duplex between the guide RNA and substrate RNA is shorter in the 3□ side of the pseudouridylation pocket, though there are some exceptions. Finally, a maximum of 2 mismatches and 3 non-canonical base pairs occur within known guide-substrate RNA interactions. These observations raise the question to the possibilities and limitations of H/ACA snoRNA interactions with potential substrate RNAs. Recently, an *in vivo* study of H/ACA guide-substrate RNA base-pairing was reported that sheds important light on the requirements for productive interactions such as the need for at least 8 base pairs between H/ACA guide and substrate RNA as well as the critical nature of base-pairs adjacent to the target uridine (De Zoysa et al. 2018). Building on these *in vivo* investigations, we aimed here at dissecting the contributions of pseudouridylation kinetics and substrate RNA affinity to the H/ACA snoRNP for efficient modification of systematically altered substrate RNAs through quantitative *in vitro* assays. Our results provide critical mechanistic information on the selection of cognate target sites by H/ACA snoRNPs as well as on the interaction of H/ACA snoRNPs with near-cognate RNA with important consequences for the cellular function of H/ACA snoRNPs.

## Results

### Base-pairing strength between substrate RNA and H/ACA snoRNPs modulates the velocity of pseudouridine formation

To investigate the impact of the H/ACA guide-substrate RNA interaction on the velocity of pseudouridine formation, short substrate RNAs were designed based on regions of yeast 25S rRNA that are complementary to the pseudouridylation pocket of the 5□ and 3□ hairpins of snR34 (designated 5□ substrate and 3□ substrate) (Fig. 1 & 2). Mismatches were introduced into the 3□ substrate by substituting nucleotides by the following rules: G to C, C to G, U to A and A to G. Adenine nucleotides were not changed to uridines to avoid the introduction of novel uridines that could potentially be pseudouridylated (Fig. 1). A short substrate RNA was also designed corresponding to a region of yeast mRNA YRA1, which had been predicted to be pseudouridylated by the 5□ hairpin of snR34 (Fig. 2) (Schwartz et al. 2014). The substrate RNAs are named according to the location within the base pairing region of the substrate RNA (5□ to 3□) in which mismatches are introduced. For example, Δ1-2, 12-17 indicates that mismatches occur at nucleotides 1-2 and 12-17 in the substrate RNA. 10CC-GG indicates that the two C nucleotides beginning at position 10 in the substrate RNA are mutated to two G nucleotides creating two GG mismatches with snR34.

**Figure 1.**
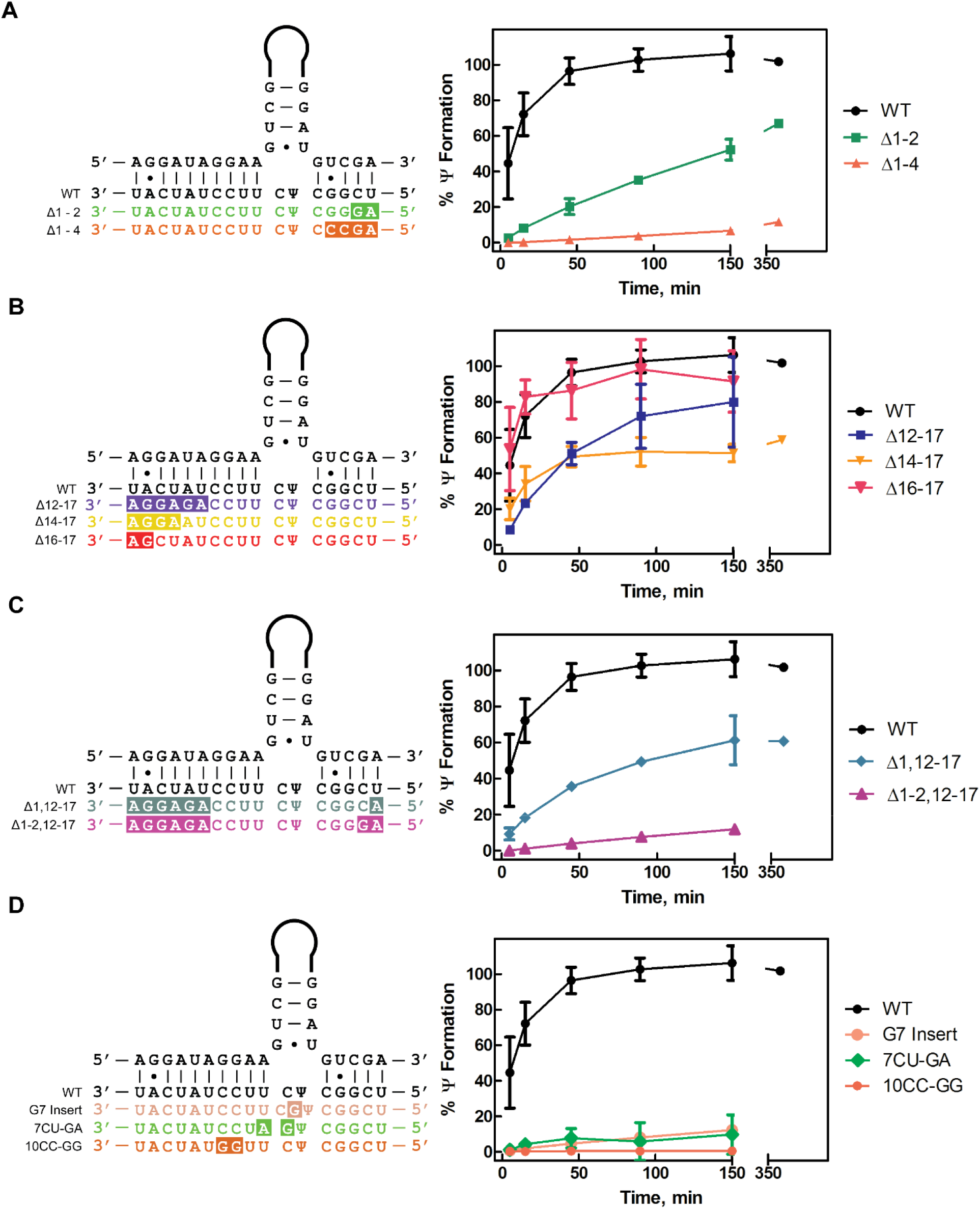
*In vitro* pseudouridylation of 3□ hairpin short substrate variants by the snR34 H/ACA snoRNP. Multiple turnover pseudouridylation reactions were performed by mixing 500 nM of each [^3^H-C5] uridine-labeled 3□ substrate RNA with 50 nM reconstituted snR34 H/ACA snoRNP. Location and sequence of substitutions are highlighted in the substrate RNA sequences on the left side, and results of the tritium release assays are shown on the right side. (A) Modification of substrate RNA variants with mismatches in the 3□ side of the pseudouridylation pocket. The wild-type substrate (black, circles), Δ1-2 substrate (green, squares) and the Δ1-4 substrate (orange, triangles) are compared. (B) Modification of substrate RNA variants with mismatches in the 5□ side of the pseudouridylation pocket. The wild-type substrate (black, circles), Δ12-17 substrate (indigo, squares), Δ14-17 substrate (yellow, inverted triangles) and Δ16-17 substrate (red, bold inverted triangles) are compared. (C) Modification of substrate variants with mismatches in both sides of the pseudouridylation pocket. The wild-type substrate (black, circles), Δ1,12-17 substrate (teal, diamonds) and the Δ1-2,12-17 substrate (magenta, triangles) are compared. (D) Modification of substrates with mismatches at internal sites in the pseudouridylation pocket. The wild-type substrate (black, circles), 10CC-GG substrate (orange, circles) 7CU-GA substrate (green, diamonds), and G7 insert substrate (peach, circles) are compared. Mean and standard deviation of three replicates are shown.

**Figure 2.**
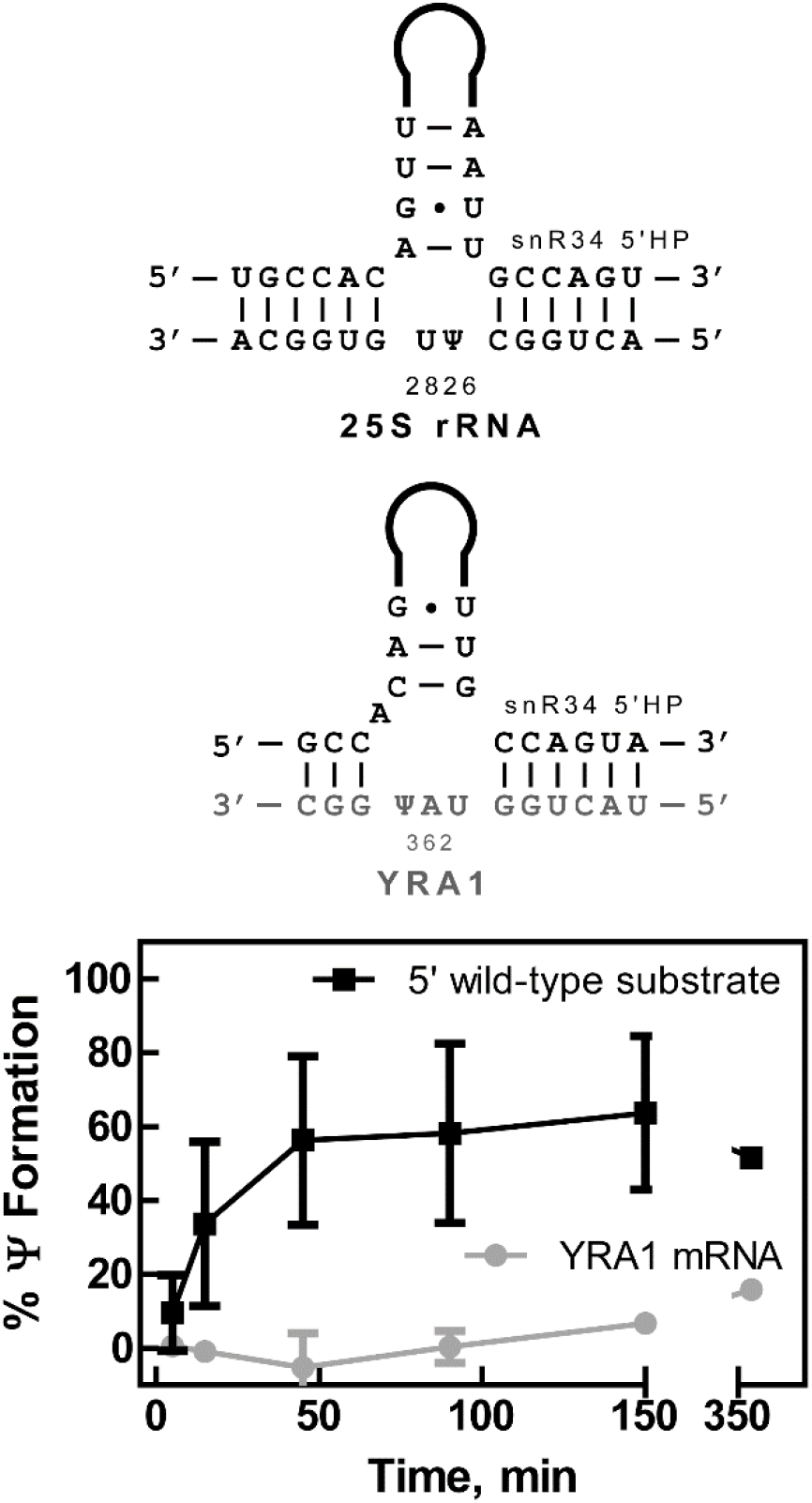
*In vitro* pseudouridylation of 5□ hairpin short substrate variants by the snR34 H/ACA snoRNP. An excess (500 nM) of each [^3^H-C5] uridine-labeled 5□ substrate RNA was incubated with 50 nM reconstituted snR34 H/ACA snoRNP. The mRNA YRA1 was predicted to be pseudouridylated by the 5□ hairpin of snR34 through the indicated base pairing (Schwartz et al. 2014). Note the difference in the snR34 hairpin base-pairing above the pseudouridylation pocket which is needed to accommodate either the wild-type (25S rRNA) substrate or the YRA1 mRNA (top panel). Pseudouridine formation in the wild-type 5□ substrate (black, squares) and in the YRA1 substrate fragment (grey, circles) is depicted in the bottom panel. Mean and standard deviation of three replicates are shown.

To quantitatively compare modification of these substrate RNAs by the snR34 H/ACA snoRNP, substrate RNAs were *in vitro* transcribed, purified and analyzed for pseudouridine formation using a tritium release assay. Substrate RNA variants that make fewer base pairs with the 3□ side of the pseudouridylation pocket (Δ1-2, and Δ1-4) display slower pseudouridine formation compared to the wild-type substrate RNA (Fig. 1A, Table 1). The wild-type substrate forms 5 base pairs with the 3□ side of the pseudouridylation pocket, whereas the Δ1-2 and Δ1-4 substrates form only 3 and 1 base pairs, respectively. The estimated initial velocity of the wild-type substrate is 26 ± 5 nM min^−1^ (Table 1). In contrast, the estimated initial velocity of the Δ1-2 and Δ1-4 substrate RNAs are only 2.0 ± 0.1 nM min^−1^ and 0.20 ± 0.02 nM min^−1^, respectively (Table 1). Thus, these substrates are modified 10- and 100-fold slower than the wild-type substrate, respectively, suggesting that reduced base pairing in the 3□ side of the pseudouridylation pocket strongly affects the rate at which pseudouridines can be formed in substrate RNAs.

**Table 1.** Activity and affinity of snR34 H/ACA snoRNP wild-type for short substrate variants. Activity was determined by tritium release assay and the initial velocity was estimated by linear regression (Fig. 1 and 2). Dissociation constants were determined by nitrocellulose filtration (Fig. 3 and S1).

| Substrate RNA | Estimated Initial Velocity<br>(nM min<sup>-1</sup>) | K<sub>D</sub> (nM) |
| --- | --- | --- |
| 3' hairpin substrate WT | 26 ± 5 | 27 ± 9 |
| 3' hairpin substrate Δ1-2 | 2.0 ± 0.1 | 26 ± 16 |
| 3' hairpin substrate Δ1-4 | 0.20 ± 0.02 | 18 ± 15 |
| 3' hairpin substrate Δ16- | 30 ± 10 | 63 ± 28 |
| 3' hairpin substrate Δ14- | 12.3 ± 0.5 | 22 ± 17 |
| 3' hairpin substrate Δ12- | 6.5 ± 0.4 | 48 ± 11 |
| 3' hairpin substrate<br>2-17 | 4.2 ± 0.5 | 24 ± 10 |
| 3' hairpin substrate Δ1-<br>-17 | 0.43 ± 0.01 | 167 ± 140 |
| 3' hairpin substrate<br>C-GG | 0.055 ± 0.03 | 266 ± 26 |
| 3' hairpin substrate 7CU- | 0.90 ± 0.2 | 76 ± 43 |
| 3' hairpin substrate G7 | $0.47 \pm 0.03$ | $51 \pm 34$ |
| 5' hairpin substrate WT | $11 \pm 0.5$ | $222 \pm 81$ |
| 5' hairpin substrate | ND | $132 \pm 28$ |

The 5□ side of the pseudouridylation pocket in the 3□ hairpin of snR34 forms ten base pairs with its wild-type substrate RNA, i.e. twice as many as the 3□ side of the pseudouridylation pocket. Therefore, we next asked whether a reduced number of base pairs on this side affects the velocity of pseudouridine formation in a similar manner (Fig. 1B). The Δ16-17 substrate RNA lacking two base pairs farthest away from the target uridine was pseudouridylated at a rate similar to the wild-type substrate (Fig. 1B, Table 1). Upon further shortening the base-pairing on the 5□ side of the pseudouridylation pocket, we observe an initial velocity of 12.3 ± 0.5 nM min^−1^ for the Δ14-17 substrate RNA and 6.5 ± 0.4 nM min^−1^ for the Δ12-17 substrate RNA (Table 1). In conclusion, removing base pairs on the 5□ side of the pseudouridylation pocket successively reduces the rate of pseudouridylation albeit not as much as removal of base pairs on the 3□ side of the pseudouridylation pocket presumably since enough base pairs are remaining, e.g. four base pairs for the Δ12-17 substrate. This bipartite base-pairing with four and five base pairs on the 5□ side and 3□ side of the pseudouridylation pocket in the Δ12-17 substrate RNA suffices to form pseudouridines with a rate that is about four-fold lower than observed for the wild-type substrate-guide RNA interaction.

After individually removing base pairs from either end of the substrate-guide RNA interaction, we then combined alterations on either side of the pseudouridylation pocket (Fig. 1C). The Δ1,12-17 substrate RNA was pseudouridylated slower than the wild type substrate, with an initial velocity of only 4.2 ± 0.5 nM min^−1^ and reaching about 60% pseudouridine formation after 150 min (Fig. 2C, Table 1). The Δ1-2,12-17 substrate RNA was modified extremely slowly with an estimated initial velocity of 0.43 ± 0.01 nM min^−1^ forming less than 20% pseudouridines after 150 min (Fig. 2C, Table 1). The extremely slow modification of the Δ1-2, 12-17 substrate RNA appears to be a combined effect of the diminished modification rates of both the Δ1-2 and the Δ12-17 substrate RNAs.

Finally, substrate RNAs that disrupt base pairs in the middle of the wild-type substrate-guide RNA interaction closer to the target uridine were analyzed. Two of these substrates introduce a bulge in the helix through mismatches with the substrate RNA (10CC-GG and 7CU-GA substrate RNA), and another substrate RNA contains an extra unpaired nucleotide adjacent to the target uridine (G7 insert). The 10CC-GG, 7CU-GA and G7 insert substrate RNAs all displayed drastically reduced initial velocities when compared to the wild-type substrate (less than 1 nM min^−1^) reaching less than 15% pseudouridine formation after 150 min (Fig. 1D, Table 1). Therefore, these substrate RNAs are not cognate, modifiable targets of the snR34 H/ACA snoRNP and are thus designated near-cognate due to their similarity to the cognate, wild-type substrate RNA.

Lastly, we tested the prediction that the snR34 5□ pseudouridylation pocket could modify the YRA1 mRNA at position 362 (Fig. 2) (Schwartz et al. 2014). Notably, binding of this mRNA would require remodelling of the pseudouridylation pocket and upper stem of the H/ACA guide RNA. We compared pseudouridylation of a wild-type 5′ substrate to the putative YRA1 mRNA substrate in tritium release assays. For the wild-type 5′ substrate, we estimated an initial velocity of 11 ± 0.5 nM min^−1^ (Table 1). In contrast, the YRA1 mRNA fragment substrate was modified at an extremely slow rate which prevented the determination of an initial velocity (Fig. 2). This result suggests that YRA1 mRNA cannot be efficiently modified by the snR34 snoRNP *in vivo*.

### H/ACA snoRNPs can bind substrate RNAs with nanomolar affinity

As the tested substrate RNAs were modified with varying reaction velocities, we asked whether the reaction velocities reflect different affinities of the snR34 H/ACA snoRNP for these substrates. Therefore, nitrocellulose filtration assays were used to determine the affinity of the snR34 H/ACA snoRNP for the different substrate RNAs. The concentration of purified Cbf5-Nop10-Gar1 does not allow titration to high concentrations. Therefore, a constant concentration of 5 nM reconstituted snR34 H/ACA snoRNP was used in the filtration experiments with increasing concentrations of radiolabelled substrate RNA as reported previously(Caton et al. 2018). Active H/ACA snoRNPs were incubated with excess substrate RNA for only 3 min such that only a low percentage of substrate RNA is pseudouridylated; therefore, we measure predominantly the binding of substrate RNA rather than product RNA to the H/ACA snoRNP. As negative control, substrate RNA titrations without H/ACA snoRNP were performed confirming a minimal background signal from substrate RNA alone.

In general, the snR34 H/ACA snoRNP binds all analyzed substrate RNAs relatively tightly with K_D_s in the nanomolar range (Fig. 3, Fig. S1, Table 1). Importantly, all measured dissociation constants are at least two-fold lower than the RNA substrate concentration (500 nM) used in pseudouridylation assays. This observation indicates that all substrate RNAs, cognate and near-cognate, can bind to the H/ACA snoRNP in the activity assay. Therefore, the lack of pseudouridine formation in the G7 insert substrate or the YRA1 mRNA fragment is not a result of insufficient binding of the near-cognate RNAs. Comparing the dissociation constants with the initial velocity of pseudouridylation also demonstrates that binding and modification are not correlated (Table 1). Reducing the number of base pairs between substrate and guide RNA successively from 15 to 8, as in the wild-type 3′ substrate compared to the Δ1,12-17 substrate RNA, does not lead to a significant change in affinity for substrate RNA. We only observe a decreased affinity when the number of base pairs is as low as seven in the Δ1-2,12-17 substrate RNA (Table 1). Of the substrates introducing substitutions at internal sites in the helices between the substrate RNA and H/ACA guide RNA pseudouridylation pocket, only 10CC-GG displays an increased dissociation constant. Otherwise, some of the near-cognate substrates that present only minimal activity are very tightly bound by the H/ACA snoRNP, such as the Δ1-4, 7CU-GA and G7 insert substrate RNAs. As a control, selected substrate RNAs were also tested for binding to catalytically inactive H/ACA snoRNPs assembled with Cbf5 D95N (Table 2, Fig. S2) which confirmed the observed trends. In conclusion, all substrate RNAs analyzed here can bind relatively tightly to the snR34 H/ACA snoRNP irrespective of their variation in base pairing to the guide RNA.

**Table 2.** Affinity of snR34 H/ACA snoRNP harboring catalytically inactive Cbf5 D95N for short substrate variants. Dissociation constants were determined by nitrocellulose filtration (Fig. S2).

| Substrate RNA | K<sub>D</sub> (nM) |
| --- | --- |
| 3' hairpin substrate WT | 232 ± 125 |
| 3' hairpin substrate Δ1,12-17 | 262 ± 57 |
| 3' hairpin substrate Δ1-2,12-17 | 542 ± 398 |

**Figure 3.**
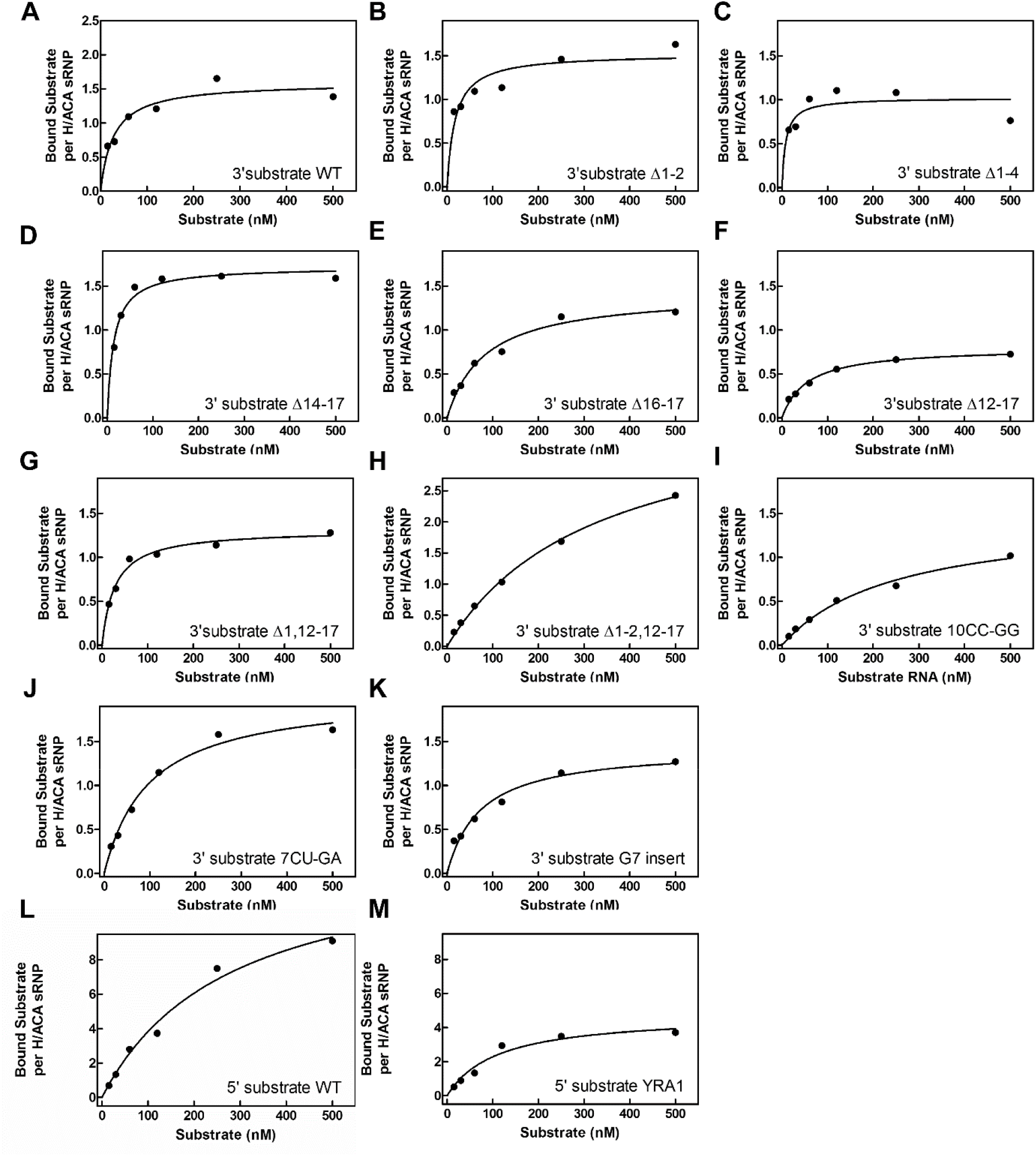
Affinity of the snR34 H/ACA snoRNP for short substrate RNA variants. Affinity of the snoRNP was determined for each substrate RNA variant using nitrocellulose filtration. Each panel shows a representative data set of triplicate experiments. Full data sets are shown in the Supplementary Figure S1. The dissociation constant (K_D_) was determined by fitting the data to a hyperbolic equation (smooth lines – see Materials and Methods). Dissociation constants are listed in Table 1. (A) 3□ hairpin substrate wild-type. (B) 3□ hairpin substrate Δ1-2. (C) 3□ hairpin substrate Δ1-4. (D) 3□ hairpin substrate Δ14-17. (E) 3□ hairpin substrate Δ16-17. (F) 3□ hairpin substrate Δ12-17. (G) 3□ hairpin substrate Δ1,12-17. (H) 3□ hairpin substrate Δ1-2,12-17. (I) 3□ hairpin substrate 10CC-GG. (J) 3□ hairpin substrate 7CU-GA. (K) 3□ hairpin substrate G7 insert. (L) 5□ hairpin substrate wild-type. (M) 5□ hairpin substrate YRA1.

### The H/ACA snoRNP can efficiently select substrates under competition

Since the H/ACA snoRNP can bind also near-cognate substrate RNAs with nanomolar affinities, but does not modify these RNAs, we asked whether the H/ACA snoRNP could select a cognate substrate RNA while in competition with a near-cognate competitor RNA. To answer this question, a competitive tritium release assay between equal concentrations of the near-cognate G7 insert RNA and the cognate wild-type 3□ substrate RNA was performed. Importantly, both substrates can form the same number and type of base pairs to snR34 (Fig. 1) and thus have comparable affinities to the snR34 H/ACA snoRNP (Table 1). Non-radiolabelled G7 insert was pre-bound to the H/ACA snoRNP before addition of a tritiated wild-type 3□ substrate RNA. Interestingly, there is no difference in the rate of pseudouridylation of the wild-type 3□ substrate RNA in the presence or absence of the competitive G7 insert sequence (Fig. 4). This observation reveals that binding of the G7 insert sequence does not competitively inhibit the binding and modification of the wild-type substrate.

**Figure 4.**
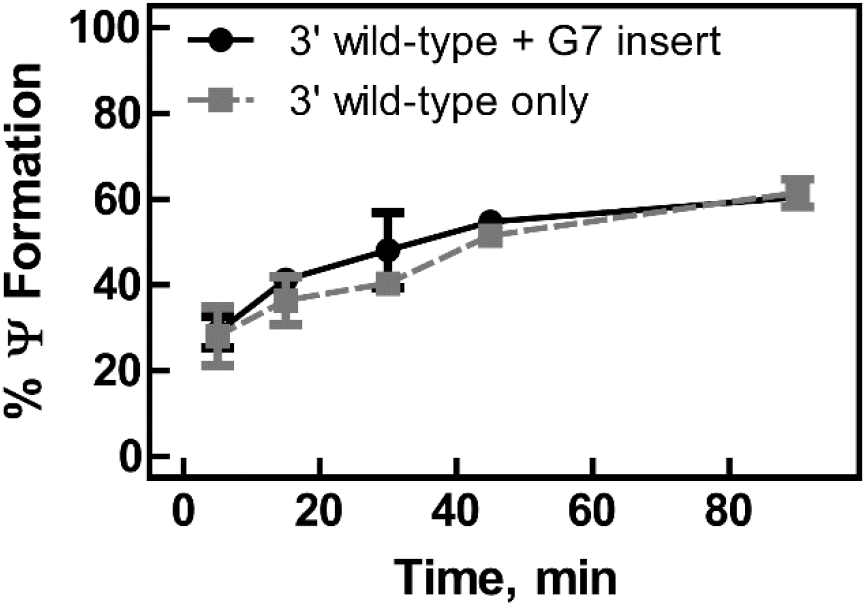
Competitive *in vitro* pseudouridylation of 3□ substrate wild-type. An excess (250 nM) of non-radioactive 3□ substrate G7 insert was incubated with 50 nM reconstituted snR34 H/ACA snoRNP for 3 min before 250 nM [^3^H-C5] uridine-labeled 3□ substrate wild-type was introduced to the reaction (black circles). As control, 250 nM [^3^H-C5] uridine-labeled 3□ substrate wild-type was incubated with 50 nM reconstituted snR34 H/ACA snoRNP in absence of 3’substrate G7 insert (grey squares). Mean and standard deviation of duplicate reactions are shown.

## Discussion

The quantitative characterization of snoRNA-substrate RNA interactions of H/ACA snoRNPs yields detailed insight into the mechanisms and consequences of RNA modification in the cell. In brief, our systematic analysis of the kinetics of pseudouridine formation by H/ACA snoRNPs with different substrate RNAs uncovers a gradual decrease of the modification velocity upon reducing the number of base pairs between substrate and guide RNA. In addition, we describe the detrimental effect of certain internal changes within the interaction between the snoRNA and its target RNA. Together with recently published data, these findings shed more light on the limitations and possibilities of targeting RNAs for pseudouridylation in the cell (De Zoysa et al. 2018). Interestingly, we demonstrate that the ability to modify a substrate RNA is not linked to the ability to bind an RNA; in other words, many near-cognate RNAs can bind with low nanomolar affinity to H/ACA snoRNPs without being pseudouridylated. This discovery provides insight into how H/ACA snoRNPs search for correct substrate RNA in the competitive cellular environment.

Knowing how pseudouridylation kinetics are modulated by substrate-guide RNA pairing has important consequences for understanding the timing and the efficiency of pseudouridine formation *in vivo*. Our results demonstrate that successively shortening the base-pairing region on either the 3′ or the 5′ side of the pseudouridylation pocket leads to a decrease in the initial velocity of pseudouridine formation, and combined reduction of base pairs on both sides further exacerbates this trend (Fig. 5). Our data also confirm that the substrate-guide RNA interaction must be bipartite with at least three base pairs on each side needed to detect more than 50% pseudouridine formation after 150 min (compare substrates Δ1-2 and Δ1-4, Fig. 1A). In addition, it has been reported that 8 base pairs are minimally required to achieve detectable pseudouridine formation in *S. cerevisiae* which is consistent with the minimal number of base pairs observed in natural snoRNA-rRNA pairs (Piekna-Przybylska et al. 2008, De Zoysa et al. 2018). In general, our findings are consistent with the reported minimal number of 8 base pairs as we only observe more than 50% pseudouridine formation after 150 min when at least 8 base pairs can be formed between substrate and guide RNA, e.g. in the Δ1,12-17 substrate. However, our data reveal a dramatic difference in the kinetics of pseudouridine formation between a natural substrate forming 15 base pairs (including a G-A base pair) and a minimal substrate with 8 base pairs since the initial velocities are about 7-fold reduced in the latter case (26 versus 4.2 nM min^−1^ for the wild-type substrate compared to the Δ1,12-17 substrate, respectively, Table 1). This finding raises the question to the *in vivo* speed and efficiency of modifying RNA. First, it is conceivable that a snoRNA, that forms more base pairs with its target rRNA, interacts with and modifies the rRNA faster than a snoRNA with less base pairs to its target. As all known H/ACA snoRNPs in yeast modify rRNA, this difference in pseudouridylation velocity could have critical implications for ribosome biogenesis (Sloan et al. 2017). Some positions in rRNA could be modified earlier than others which could affect rRNA folding and protein association. Second, different velocities of rRNA modification by H/ACA snoRNPs suggests that there might not be enough time to ensure complete pseudouridine formation at all target sites as the rRNA folds into the compact ribosome structure. Recent findings suggest that not all rRNA sites are stoichiometrically modified *in vivo* which could result in ribosome heterogeneity with potential functional consequences for the translation of selected mRNAs (Henras et al. 2017). Indeed, reduced pseudouridine content in ribosomes of Dyskeratosis congenita patients harboring mutations in the DKC1 gene (the homolog of yeast Cbf5) has been associated with effects on internal ribosome entry site (IRES) mediated translation (Yoon et al. 2006, Penzo et al. 2015). It is also known that the abundance of certain snoRNAs varies between tissues, which could in turn influence the velocity and efficiency of rRNA pseudouridylation (McMahon et al. 2015). In conclusion, our discovery that base pairing numbers influence the kinetics of pseudouridine formation by H/ACA snoRNPs is important for understanding the timing and the efficiency of rRNA modification during ribosome biogenesis with consequences for ribosome functionality.

**Figure 5.**
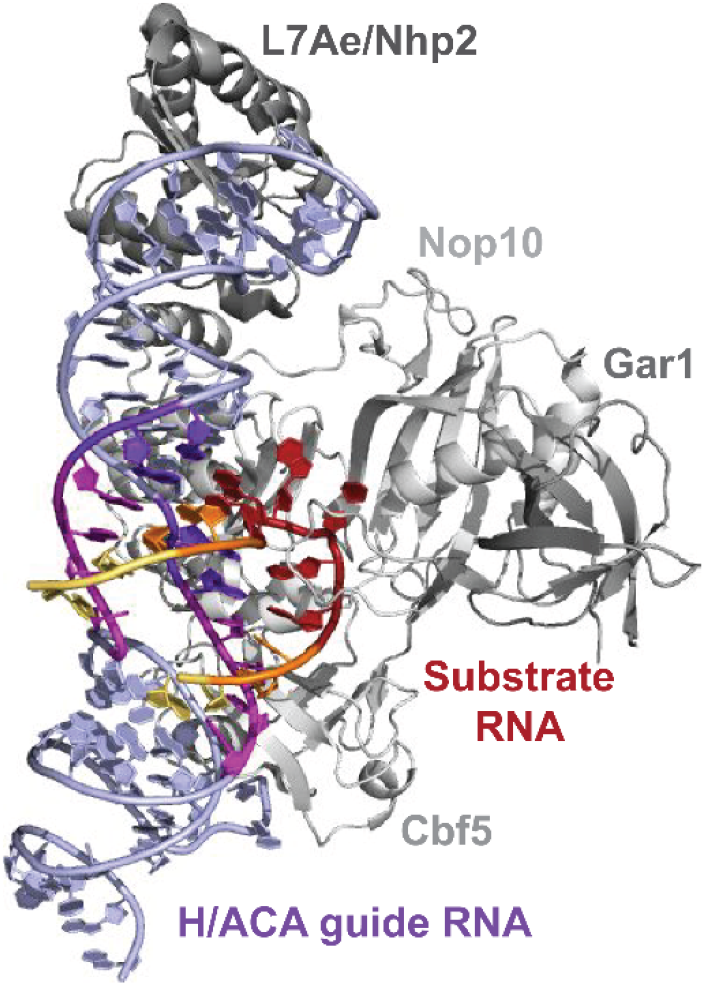
H/ACA snoRNP structure highlighting the important interactions between H/ACA guide RNA and substrate RNA. Ribbon structure of an archaeal H/ACA snoRNP complex assembled on a single-hairpin H/ACA guide RNA with a short substrate RNA bound (PDB ID: 3HAY). The proteins are depicted and labelled in grey whereas the guide RNA is shown in blue to purple and the substrate RNA in red to yellow. The first two base-pairing interactions on either side of the target uridine are essential for pseudouridylation and are colored red and dark blue. The next two important base-pairing interactions are highlighted in orange and purple. The less important base pairs at the bottom of the pseudouridylation pocket are shown in yellow and pink.

In addition to considering the total number of (continuous) base pairs between substrate and guide RNA, it is arguably even more important to understand the complex effects of internal mismatches between the substrate and guide RNA as many such imperfect sequences will be encountered by H/ACA snoRNPs in the cell. Here, we tested three different substrate RNAs with changes close to the target uridine which all dramatically reduced the initial velocity of pseudouridine formation (Fig. 1, Table 1). For the 10CC-GG substrate, the loss of pseudouridine formation could result from the reduced affinity of substrate RNA binding to the H/ACA snoRNP (Table 1). In addition, the 10CC-GG substrate could be unable to stably position the target uridine in the active site of Cbf5 as the lack of the two C-G pairs could further destabilize the adjacent A-U pairs flanking the target uridine due to loss of stacking interactions. Similarly, the 7CU-GA substrate is lacking a base pair in the 5′ side of the pseudouridylation pocket that is directly flanking the unpaired dinucleotide including the target uridine. As shown previously, this base pair is particularly important for pseudouridine formation (De Zoysa et al. 2018). Interestingly, in our case, the insertion of an additional unpaired nucleotide adjacent to the target uridine (G7 insert substrate) also abolished pseudouridine formation; presumably, the steric crowding of three nucleotides prevents the target uridine from entering the catalytic site. This finding stands in contrast to published data that up to four additional uridines can be accommodated next to the target uridine in the 5′ hairpin of snR81 (De Zoysa et al. 2018). Further studies are needed to resolve this discrepancy. For example, it is conceivable that pyrimidines are more easily accommodated next to the target uridine than purines. Lastly, our detailed analysis of substrate-guide RNA interactions also reveals that predictions for active combinations *in vivo* must be made very carefully. Clearly, our results show that the YRA1 mRNA cannot be modified by snR34 as suggested (Schwartz et al. 2014) (Fig. 2). This observation can be explained by the fact that three unpaired nucleotides including a purine would have to be accommodated for this substrate-guide RNA combination, and furthermore the target uridine would be on the wrong side of these three nucleotides; both features likely render the substrate not suitable for modification by the snR34 H/ACA snoRNP. In conclusion, our data indicate that the removal of base pairs distant from the target uridine leads only to a gradual reduction on the rate of pseudouridylation whereas changes or insertions of nucleotides close to the target uridine typically cannot be tolerated for pseudouridylation by H/ACA snoRNPs.

Interestingly, most substrate RNAs tested in our study can bind to the snR34 H/ACA snoRNP with a high affinity, i.e. a dissociation constant under 100 nM (Fig. 3, Table 1). Thus, the number of base pairs ranging from 15 to 8 does not significantly influence the affinity of the H/ACA snoRNP for substrate RNA. Based on structural studies, the substrate RNA binds in an omega conformation to H/ACA guide RNA, and the base-pairing on the 5′ side of the pseudouridylation pocket is stabilized through stacking with the upper stem of the H/ACA guide RNA whereas the base pairing on the 3′ side of the pseudouridylation pocket is supported by direct interactions with Cbf5 (Fig. 5) (Liang et al. 2007, Duan et al. 2009). If a minimal number of three to four base pairs is formed on either side of the pseudouridylation pocket, these interactions result in an overall tight binding of substrate irrespective of the exact number of base pairs. It is also conceivable that the omega conformation of the substrate RNA generates increasing strain in the substrate RNA as more base pairs are formed. Accordingly, the base pairs at the bottom of the pseudouridylation pocket might be less stable and might fluctuate between a base-paired and non-base-paired state. Consequently, the base pairs at the top of the pseudouridylation pocket, that are common among all active substrate RNAs, would predominantly contribute to the affinity of substrate RNA explaining the observed similar affinities.

Notably, even substrate RNAs barely modified by the snR34 H/ACA snoRNP are bound with relatively high affinity by the H/ACA snoRNP (e.g. the 7CU-GA and G7 insert substrate RNAs, as well as the YRA1 mRNA fragment). First, this finding clearly rules out impaired binding as the cause for lack of activity for these near-cognate substrates. Thus, the only conceivable explanation for the absence of pseudouridine formation is that these substrates bind incorrectly without positioning the target uridine in the active site of Cbf5. Second, this finding raises several questions on the interactions of H/ACA snoRNPs with other near-cognate RNAs in the cell. Obviously, H/ACA snoRNPs can bind many imperfectly base pairing, near-cognate RNAs without modifying these. However, it is important that H/ACA snoRNPs do not remain bound to near-cognate RNAs too long as this would inhibit their activity. Indeed, we have shown that wild-type substrate efficiently competes with the inactive G7 insert substrate which forms the same base pairs to snR34 guide RNA (Figs. 1 and 4). This finding can be explained if the near-cognate G7 insert substrate dissociates rapidly from the H/ACA snoRNPs. In the cell, H/ACA snoRNPs are therefore likely to rapidly bind and dissociate from near-cognate RNAs as a mechanism to search for the correct target site among a large range of RNA sequences. Further studies are required to fully elucidate the substrate screening mechanism of H/ACA guide RNAs, but it is conceivable that the two pseudouridylation pockets within the H/ACA guide RNAs could alternate in binding to near-cognate RNA thereby keeping the H/ACA snoRNP near rRNA in the nucleolus. Moreover, it has been speculated that H/ACA snoRNPs could act as rRNA chaperones during the early stages of ribosome biogenesis by unfolding the pre-rRNA (Sloan et al. 2017). The transient binding of H/ACA snoRNPs to near-cognate sites in rRNA could further contribute to this potential chaperone function of H/ACA snoRNPs as they could keep these near-cognate sites in rRNA unfolded in addition to their cognate target sites.

In summary, our study provides critical insight into the selection of target sites for pseudouridylation by H/ACA snoRNPs. Our findings on differential kinetics of pseudouridine formation as well as transient binding to near-cognate sites have implications for the mechanism, timing and efficiency of rRNA modification and ultimately ribosome function. In the future, it will be interesting to further investigate the kinetics of RNA association and dissociation of H/ACA snoRNPs and their impact on rRNA structure. In addition, the results presented here will allow us to better predict new, active substrate-guide RNA pairings, for example for the many orphan H/ACA guide RNAs in humans or for the many pseudouridine sites discovered in human and yeast mRNAs and non-coding RNAs that could be formed by H/ACA snoRNPs.

## Materials & Methods

### Reagents

[5-^3^H] UTP for *in vitro* transcriptions was purchased from Moravek Biochemicals, and [γ-^32^P] ATP for guide RNA 5′ end labelling was obtained from Perkin Elmer. DNA oligonucleotides were obtained from Integrated DNA Technologies (IDT). Polymerase chain reaction (PCR) reagents and *Pfu* DNA polymerase were purchased from Truin Science. All other chemicals were purchased from Fisher Scientific.

### *In vitro* transcription, and purification of substrate RNA variants

The gene encoding the H/ACA guide RNA snR34 was PCR-amplified to include a T7 promoter as previously described (Caton et al. 2018). Template DNA for substrate RNA was generated from PCR extension of two partially overlapping oligonucleotides (Table 3) (Milligan et al. 1987). Radioactive substrate RNAs were generated by *in vitro* transcriptions including 3 mM ATP, CTP and GTP, and 0.1 mM [5-^3^H] UTP (16.2 Ci/mmol). RNAs were purified by crush and soak gel extraction from a 15% urea-polyacrylamide gel. The RNA band was identified by UV shadowing, and the gel area was excised, crushed, and incubated in 1x TBE for 6 h. After centrifugation, and phenol-chloroform extraction, the RNA was ethanol precipitated, resuspended in deionized water and stored at −20 °C. RNA concentration was determined by A_260_ using extinction coefficients calculated by OligoAnalyzer 3.1 (IDT), and the specific activity was determined by scintillation counting.

**Table 3.**
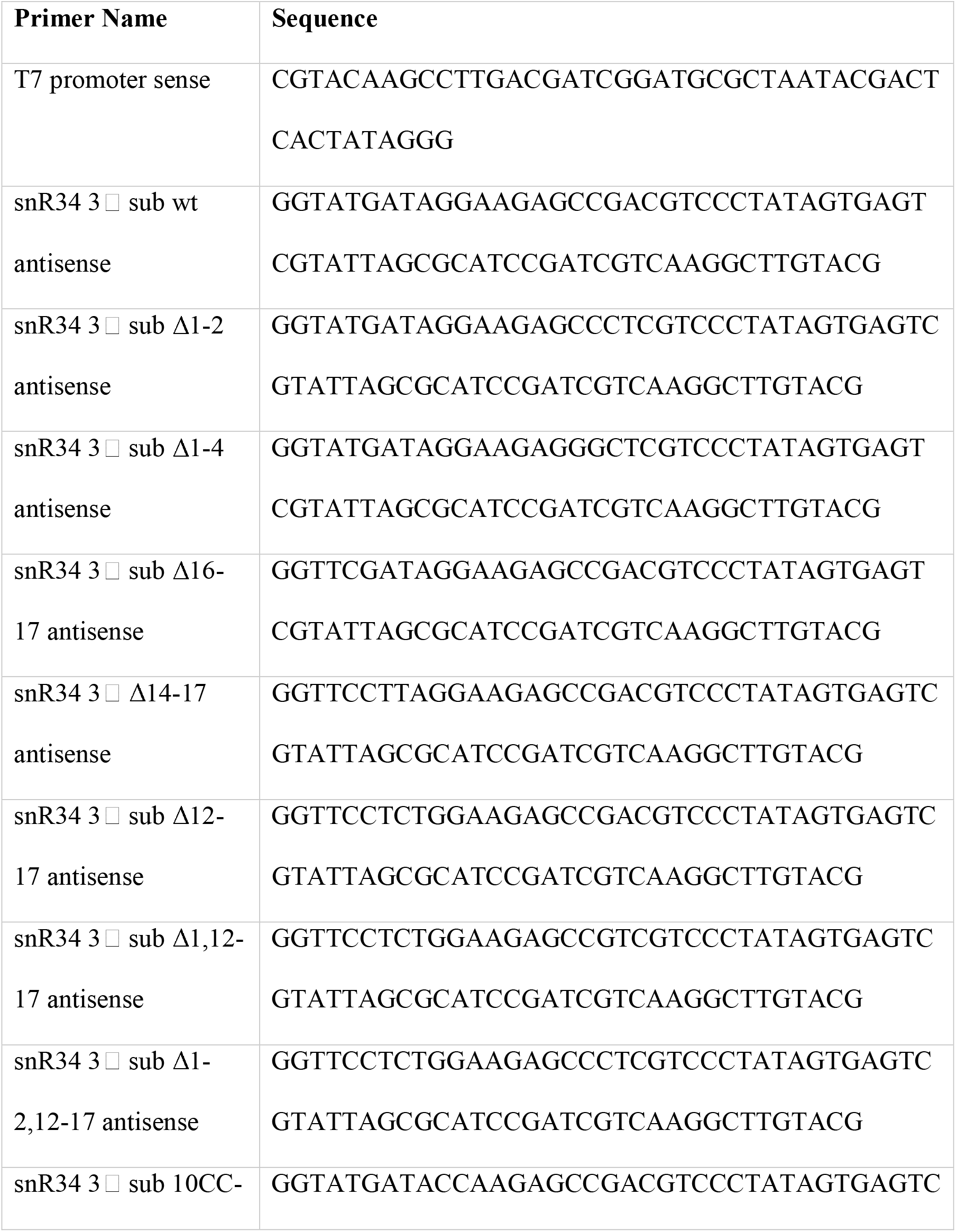
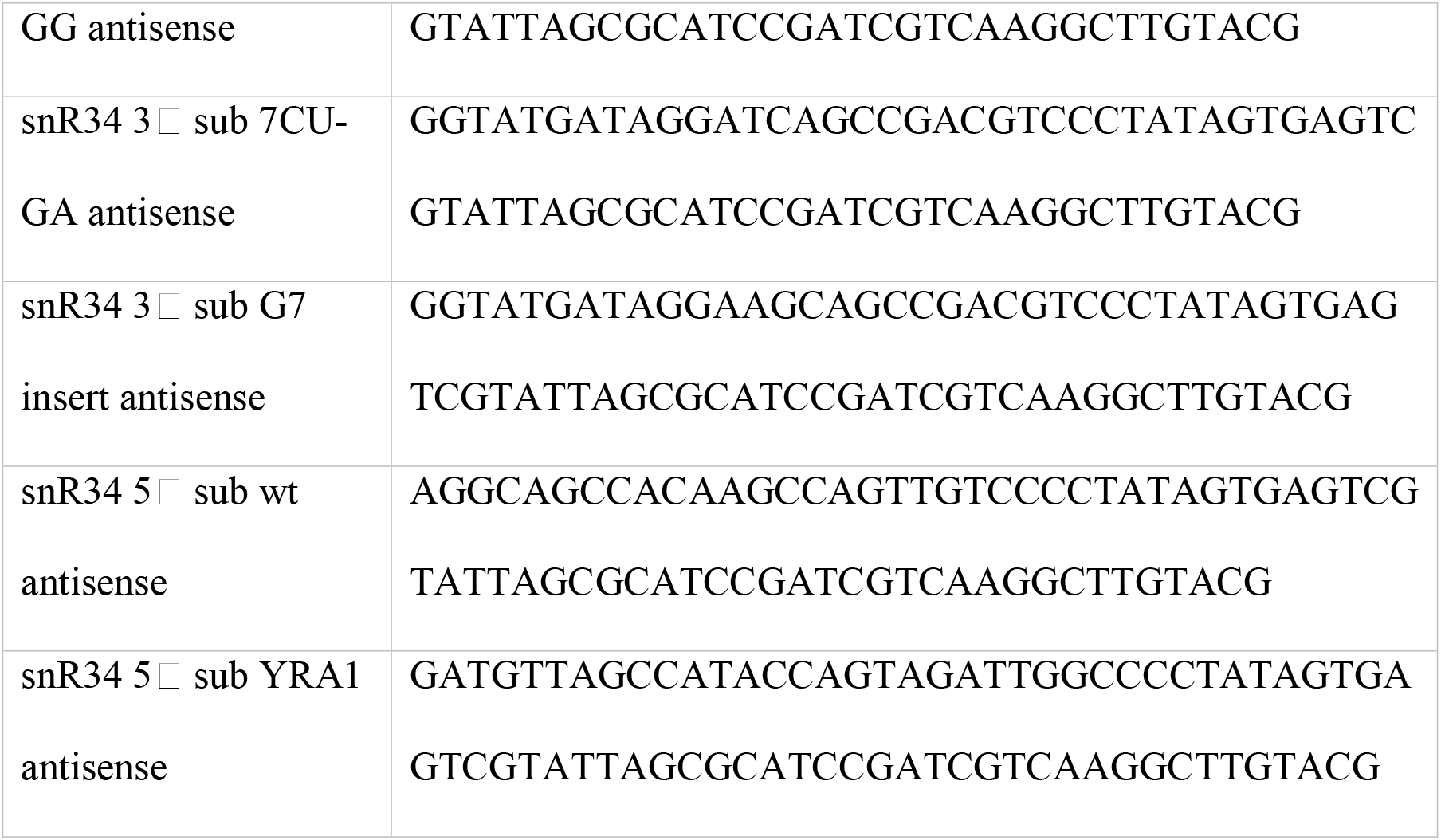
Oligonucleotides for *in vitro* transcription template generation for snR34 5□ and 3□ substrates. All oligos are shown in the 5□ to 3□ direction.

### Reconstitution of H/ACA snoRNPs

The protein Nhp2 and the complex of Cbf5(wt or D95N)-Nop10-Gar1 was recombinantly overexpressed in *Escherichia coli* and purified as described (Caton et al. 2018). Full length snR34 was refolded by heating to 75°C for 5 min and cooling slowly to room temperature. Guide RNA was combined with Cbf5, Nop10, Gar1 and Nhp2 in a 0.45:1 guide RNA: protein ratio in Reaction Buffer (20 mM HEPES-KOH pH 7.4, 150 mM NaCl, 0.1 mM ethylenediaminetetraacetic acid (EDTA), 1.5 mM MgCl_2_, 10 % (v/v) glycerol, 0.75 mM DTT). The mixture was incubated for 10 min at 30 °C to allow complex formation.

### Tritium Release Assay

Multiple turnover assays were performed with 50 nM reconstituted H/ACA snoRNP and 500 nM substrate RNA. The modification reaction was performed at 30 °C. Samples containing 7.5 – 25 pmol of RNA (depending on the specific activity) were taken, quenched in 1 mL 5 % (w/v) activated charcoal (Norit A) in 0.1 M HCl. After centrifugation, 850 μL of the supernatant was mixed with 300 μL 5% Norit A (w/v) in 0.1 M HCl and centrifuged again. The supernatant was filtered through glass wool and 800 μL of the filtrate was subjected to scintillation counting to determine the amount of tritium released corresponding to the amount of pseudouridine formed. Data was analyzed using GraphPad Prism (La Jolla, California, USA). Initial velocities were estimated by linear regression of the initial region of the tritium release assay time course (<70% of the measured end-level) and forcing the fitted line through zero. As the substrate concentration is relatively high with 500 nM (at least two-fold higher than the K_D_ for substrate binding), it is reasonable to assume that we are measuring significantly above the K_M_. Hence, the observed initial velocity will be close to the maximal reaction velocity v_max_, where the initial velocity is relatively insensitive to small variations in the substrate concentration. This allowed us to determine the initial velocities from time points where less than 70 % substrate has been converted to product.

### Nitrocellulose Filtration Assay

Binding assays using tritium-labelled substrate RNAs were performed by incubating increasing concentrations of substrate in the presence of 5 nM H/ACA snoRNPs reconstituted with Cbf5 wild-type or the inactive D95N variant for 3 min at 30 °C. The complete 200 μL reaction was filtered through a nitrocellulose membrane, followed by washing of the nitrocellulose membrane with 1 mL cold Reaction Buffer. The nitrocellulose membrane was dissolved in 10 mL EcoLite scintillation cocktail (EcoLite (+), MP Biomedical) followed by scintillation counting to determine the amount of substrate RNA bound to the H/ACA snoRNP.

Dissociation constants (*K*_D_) were determined by fitting the binding curves to the hyperbolic function in GraphPad Prism

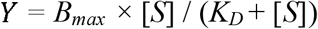

where [S] is the substrate concentration and B_max_ is the maximum binding. The substrate RNA: enzyme ratio was calculated by dividing the picomoles of substrate RNA retained on the nitrocellulose membrane by the picomoles of enzyme in the reaction.

## Acknowledgements

We thank Jane Jackman for valuable feedback on this study. This work was supported by Alberta Innovates [Strategic Research Chair 2015]; the University of Lethbridge [ULRF 2014, HRAF 2015]. E.K.K. received an NSERC CGS-M scholarship and a Queen Elizabeth II graduate scholarship. Funding for open access charge: Alberta Innovates.

---

**Figure S1.**
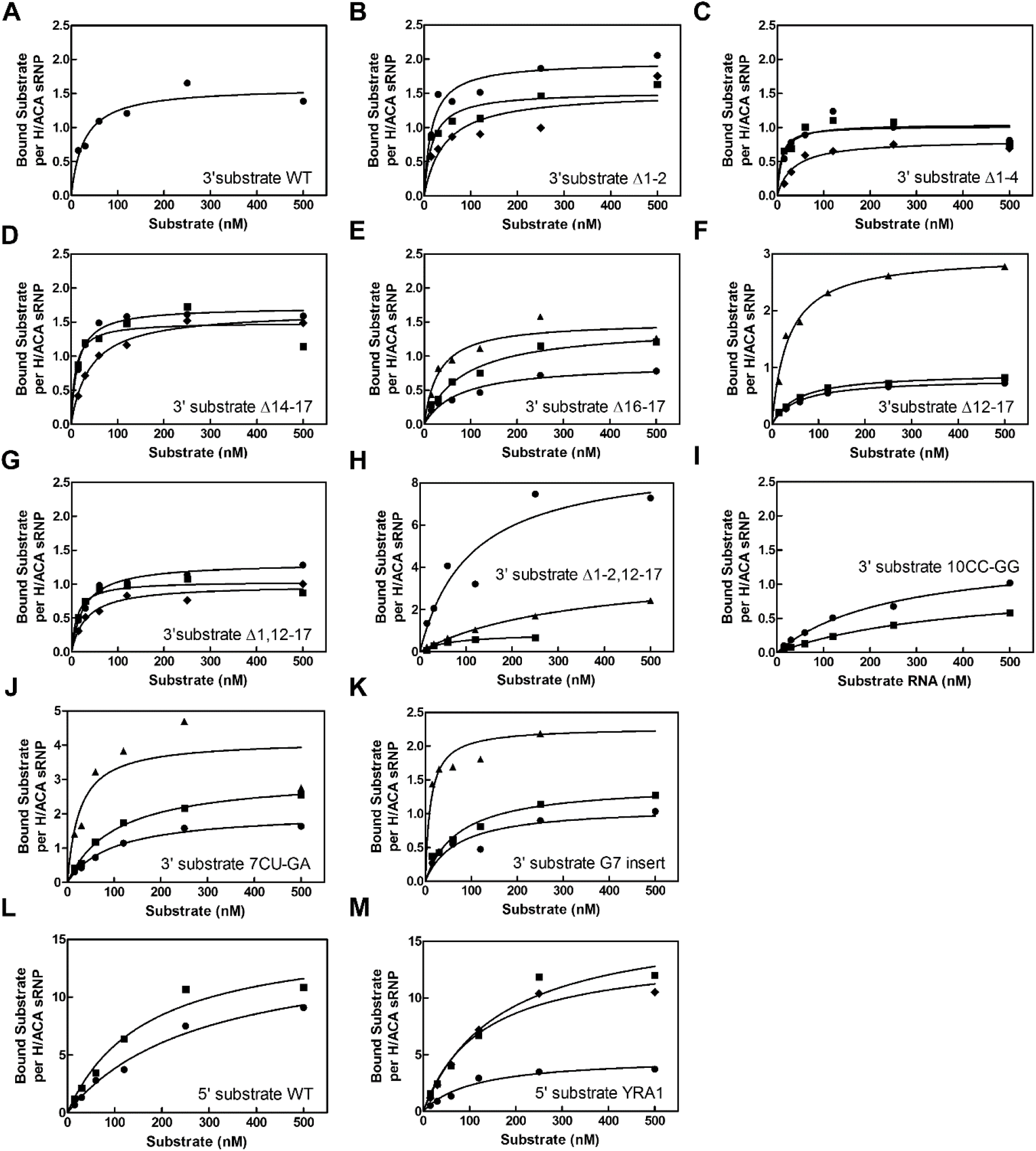
Affinity of the snR34 H/ACA snoRNP for short substrate RNA variants. Affinity of the wild-type snoRNP was determined for each substrate RNA variant using nitrocellulose filtration. The dissociation constant (K_D_) was determined by fitting the data to a hyperbolic equation (smooth lines – see Materials and Methods). Dissociation constants are listed in Table 4. (A) 3□ hairpin substrate wild type (previously determined by (Caton, Kelly et al. 2018)). (B) 3□ hairpin substrate Δ1-2. (C) 3□ hairpin substrate Δ1-4. (D) 3□ hairpin substrate Δ14-17. (E) 3□ hairpin substrate Δ16-17. (F) 3□ hairpin substrate Δ12-17. (G) 3□ hairpin substrate Δ1,12-17. (H) 3□ hairpin substrate Δ1-2,12-17. (I) 3□ hairpin substrate 10CC-GG. (J) 3□ hairpin substrate 7CU-GA. (K) 3□ hairpin substrate G7 insert. (L) 5□ hairpin substrate wild type. (M) 5□ hairpin substrate YRA1.

**Figure S2.**
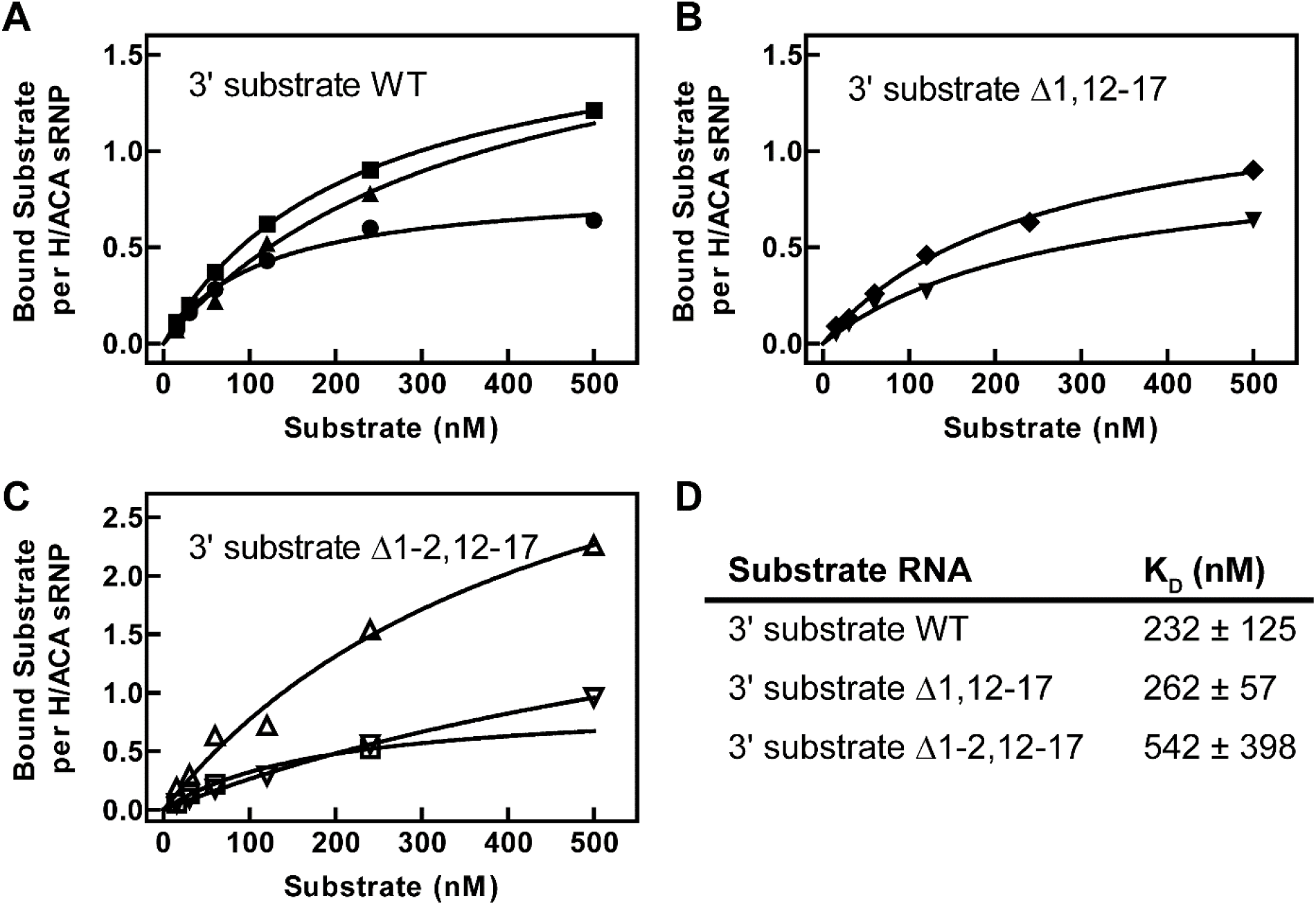
Affinity of the snR34 H/ACA snoRNP containing the inactive Cbf5 D95N variant for selected short substrate RNA variants. Affinity of the Cbf5 D95N snR34 snoRNP was determined for each substrate RNA variant using nitrocellulose filtration. (A) 3□ hairpin substrate wild type. (B) 3□ hairpin substrate Δ1,12-17. (C) 3□ hairpin substrate Δ1-2,12-17. (D) Summary of dissociation constants of substrate RNAs to the catalytically inactive Cbf5 D95N snR34 snoRNP.

